# Symbiotic root infections in *Medicago truncatula* require remorin-mediated receptor stabilization in membrane nanodomains

**DOI:** 10.1101/179036

**Authors:** Pengbo Liang, Thomas F. Stratil, Claudia Popp, Macarena Marín, Jessica Folgmann, Kirankumar S. Mysore, Jiangqi Wen, Thomas Ott

## Abstract

Plant cell infection is tightly controlled by cell surface receptor-like kinases (RLKs) Alike other RLKs the *Medicago truncatula* entry receptor LYK3 laterally segregates into membrane nanodomains in a stimulus-dependent manner. Although nanodomain localization arises as a generic feature of plant membrane proteins, molecular mechanisms underlying such dynamic transitions and their functional relevance remained poorly understood. Here, we demonstrate that actin and the flotillin protein FLOT4 form the primary and indispensable core of a specific nanodomain. Infection-dependent induction of the remorin protein and secondary molecular scaffold SYMREM1 results in subsequent recruitment of ligand-activated LYK3 and its stabilization within these membrane subcompartments. Reciprocally, the majority of this LYK3 receptor pool is destabilized at the plasma membrane and undergoes rapid endocytosis in *symrem1* mutants upon rhizobial inoculation resulting in premature abortion of host cell infections. These data reveal that receptor recruitment into nanodomains is indispensable for their function during host cell infection.

**SIGNIFICANCE STATEMENT:** Pattern recognition receptors control the cellular entry of pathogenic as well as symbiotic microbes. While ligand-induced changes in receptor mobility at the plasma membrane and their localization in membrane nanodomains appears as a general feature, the molecular mechanism and the biological relevance of this phenomenon remained unknown. Here, we show that immobilization of the symbiotic cell entry receptor LYK3 in nanodomains requires the presence of actin and the two molecular scaffold proteins FLOT4 and SYMREM1. While FLOT4 forms the initial core structure, infection-induced expression and subsequent physical interaction of SYMREM1 with LYK3 stabilizes the activated receptors in membrane nanodomains. This recruitment prevents its stimulus-dependent endocytosis and ensures progression of the primary infection thread into root cortical cells.

## INTRODUCTION

Physical recruitment of membrane-resident ligand-binding receptors into membrane nanodomains emerges as a common phenomenon within the plant kingdom (1, 2). It represents a possible generic theme for the activation and maintenance of receptor-controlled signal transduction. Indeed, a number of plant receptor-like kinases (RLKs), as well as receptor-associated and immunity-related proteins localize to nanodomains (1-3). Given the diversity of co-existing nanodomains (4), these membrane compartments might serve as active hubs that allow a spatially confined assembly of pathway-specific and functionally related signalling complexes (3). The fact that multivalent and nanodomain-localized molecular scaffold proteins associate and/or co-localize with RLKs and other signalling proteins (5-8) makes them potent candidates to act as central integrators of these membrane subcompartments. In plants, PM-resident scaffolding functions have been proposed for flotillins and remorins (9). These soluble proteins associate with the inner leaflet of the PM and interact with different partners (6, 8, 10-15). There, they contribute to phytohormone signalling (5), developmental processes (13, 16), abiotic stress responses (17, 18), plant-virus interactions (14, 15, 19, 20) and root nodule symbiosis (6, 7, 11). Latter mutualistic interaction between rhizobia and legumes is initiated at the epidermis by a highly specific molecular dialogue between the two partners (21). In *Medicago truncatula* the primary perception of rhizobial lipo-chitooligosaccharides, the so-called ‘Nod Factors’ (NF), and subsequent colonization of host root hair cells by symbiotic bacteria is tightly controlled by LysM-type RLKs such as the ligand binding receptor NOD FACTOR PERCEPTION (NFP) and the co-receptor LYSINE MOTIF KINASE 3 (LYK3) (22, 23). Rhizobial infection is preceded by NF- and microbe-triggered changes in root hair morphology (root hair deformation and root hair curling) (24). This results in an inclusion of rhizobia and formation of an infection chamber inside the root hair curl, which is followed by an invagination of the host cell plasma membrane and the subsequent formation of an infection thread (IT) (25). Molecularly, infection-induced activation of LYK3 results in a transition of its membrane-partitioning including restricted lateral mobility and accumulation in FLOTILLIN 4 (FLOT4)-labelled nanodomains (26). Besides FLOT4, two other scaffolds, FLOT2 and the remorin SYMREM1, contribute to host-driven infection control (6, 11) in a so far elusive molecular manner. Here, we chose rhizobial infections as an appropriate biological system to unravel the spatio-temporal sequence of molecular events that control protein patterning at the host cell surface. Furthermore, we genetically tested whether receptor recruitment into nanodomains is functionally relevant at all.

## RESULTS AND DISCUSSION

In order to precisely define the experimental setup, we first determined whether SYMREM1 and FLOT4 function in the same pathway during rhizobial infections of *M. truncatula* root hairs. Since stringent control over infection requires tightly coordinated cell-type-specific regulatory networks, two transcriptional reporters expressing nuclear-targeted tandem GFPs (*ProSYMREM1-NLS-2xGFP* and *ProFLOT4-NLS-2xGFP*) were independently transformed into *M. truncatula* wild-type roots. For *SYMREM1* no nuclear reporter activity was observed in the absence of *S. meliloti* (Fig. 1A) while nuclear fluorescence was clearly detected in curling as well as adjacent trichoblasts and atrichoblasts one day after rhizobial inoculation (Fig. 1B-C’). In contrast, the *FLOT4* promoter was constitutively active at low levels in all epidermal cells of the infection zone even in the absence of rhizobia (Fig. 1D) and was further induced upon inoculation (Fig. 1E-F’). To precisely localize the encoded proteins during root hair infection, we independently expressed FLOT4 and SYMREM1 as GFP fusion proteins under the control of their respective endogenous promoters in the A17 wild-type background. Due to low signal intensities, GFP fluorescence was enhanced by immune-staining using Atto488-coupled anti-GFP-nanobodies. Both scaffold proteins localized to IT membranes and accumulated around the primary infection thread and around the infection chamber in curled root hairs seven days post inoculation (dpi) with *Sinorhizobium meliloti* (Fig. 1 G-H’). This resembles the localization pattern of LYK3 when being expressed as a GFP-fusion protein driven by the LYK3-promoter (Fig. 1 I, I’) and those previously published (26).

**Fig. 1.**
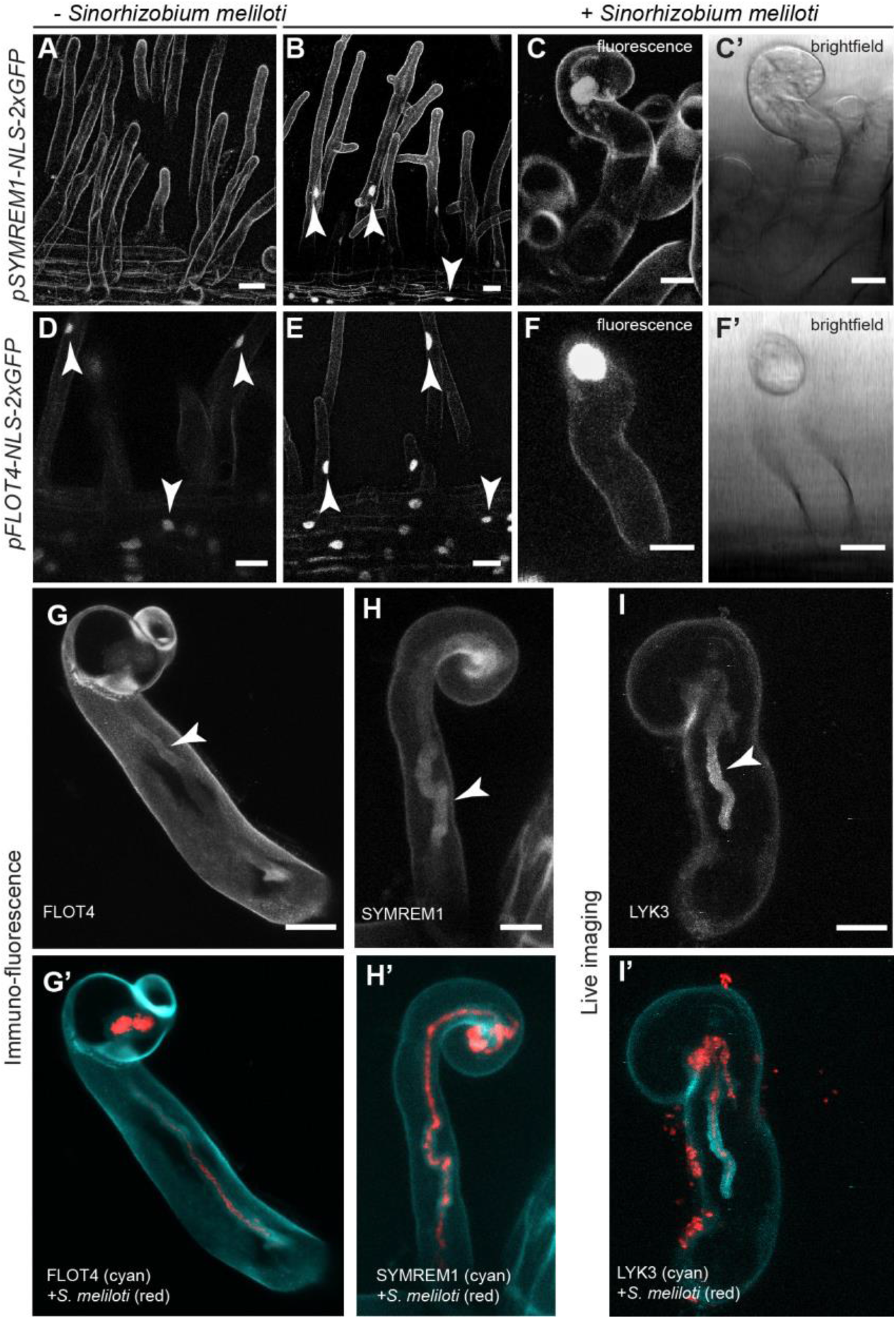
Expression and localization analyses of *SYMREM1* and *FLOT4*. Monitoring promoter activation of *SYMREM1* (*A-C’*) and *FLOT4* (*D-F*’) with cellular resolution in transgenic *M. truncatula* root hairs by using the genetically encoded reporters ProSYMREM1:NLS:2xGFP and ProFLOT4:NLS:2xGFP, respectively. Activation of *SYMREM1* as indicated by nuclear fluorescence (arrowheads) in uninoculated (*A*) and *S. meliloti* 2011 (mCherry) inoculated (*B*) roots. (*C-C’, F-F’*) Close-ups of individual curled root hairs with fluorescent nuclei. (*A-C’’*). Autofluorescent contours of roots hairs and epidermal cells are visible due to ultra-sensitive imaging settings that were chosen due to low signal intensities. (*D-F’*) Activation of the *FLOT4* promoter in uninoculated (*D*) and inoculated (*E*) conditions (arrowheads). (*G-H’*) Immuno-fluorescence targeting FLOT4-GFP (*G, G’*) and GFP-SYMREM1 (*H, H’*), both expressed from their endogenous promoters. (*I, I’*) Transgenic line expressing a ProLYK3::LYK3-GFP construct. Arrowheads indicate the localization of the respective protein around the ITs. Scale bars indicate 20 μm in *A-B and D-E* and 10 μm in *C-C’, F-F’, G, H, I.*

So far, the importance of flotillins and SYMREM1 during primary root hair infection had only been assessed by RNAi-mediated knock-down in transgenic roots (6, 11). To test overlapping functions genetically we isolated three novel independent *flot4* mutant alleles: NF14593 (*flot4-1*), NF4565 (*flot4-2*), NF14107 *flot4-3*) (Fig. 2A) by PCR screening of the *Tnt1* transposon insertion population at the Noble Foundation (27). While we were unable to recover homozygous offspring of the *flot4-1* allele, transcript levels were not significantly decreased in homozygous T2 *flot4-2* plants. However, *FLOT4* mRNA was barely detectable in *flot4-3* mutant plants confirming that it can be considered a transcriptional null-allele (Fig. S1A). For *FLOT2,* two independent lines were obtained (Fig. 2A). However, despite PCR-based screening of 64 and 95 individuals for *flot2-1* (NF19805) and *flot2-2* (NF13963), respectively, no homozygous T2 plants were recovered indicating lethality of this mutation. Similarly, such phenotype was also reported for AtFLOT1, the closest homolog of MtFLOT2 from *A. thaliana* (13). Consequently, we had to exclude FLOT2 from further analysis. In addition to a previously identified *symrem1* insertion line (NF4432; *symrem1-1*) (6) we also identified a second mutant allele (NF3495; *symrem1-2*) (Fig. 2A). *SYMREM1* transcripts were significantly reduced in both alleles when plants were inoculated for 24 hours with *S. meliloti* (Fig. S1B). Next, we phenotypically characterised these mutants by assessing primary infection success of rhizobia at 7 dpi. For this, we scored infected root hairs containing micro-colonies (MCs)/aborted infections or elongated infection threads reaching into cortical cells. In the *flot4-3* (n=14), *symrem1-1* (n=18) and *symrem1-2* (n=18) mutants rhizobia were trapped inside of curled root hairs mostly at the microcolony in 60%, 76% and 67% of the cases, respectively (Fig. 2B, E, F). Consequently, only 40% (flot4-3), 24% (*symrem1-1*) and 33% (*symrem1-2*) of all elongated ITs passed the epidermis/cortical cell boundary, while this was observed in 78% of all infection in the WT (Fig. 2C,D).

**Fig. 2.**
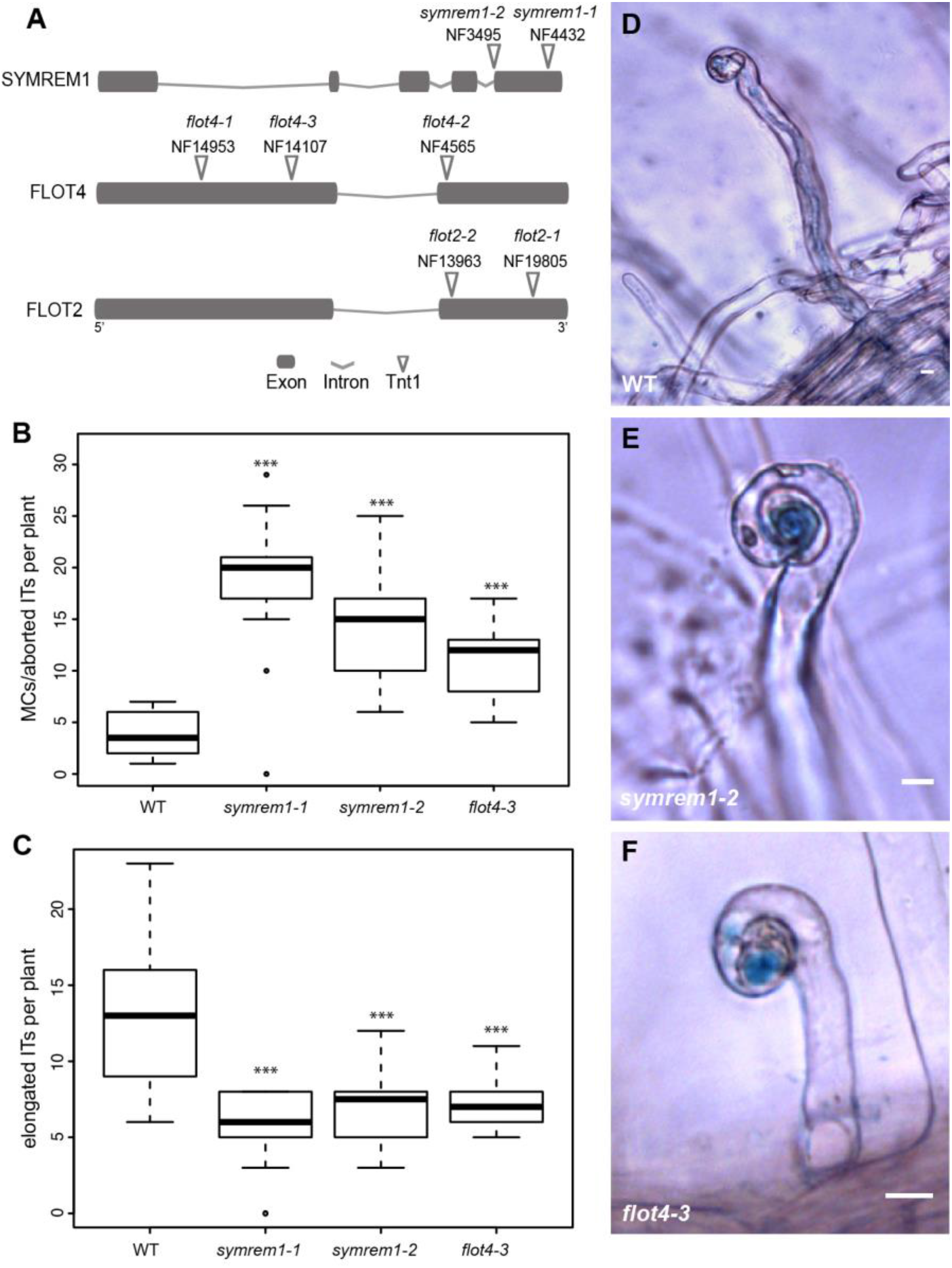
*symrem1* and *flot4* mutants are impaired in infection thread initiation. (*A*) Mapped *Tnt1* transposon insertions. Phenotypical analysis of *symrem1* and *flot4* mutants. Infection structures were scored 7 days after inoculation with *S. meliloti* (lacZ) and classified into micro-colony (MC)/aborted infection threads (IT) that were arrested within trichoblasts (*B*) and elongated ITs that progressed into the root cortex (*C*). Representative elongated ITs as found in WT (*D*), and infections arrested in the MC stage in *symrem1-2* (*E*) and *flot4-3* (*F*) mutants. (*B-C*) Asterisks indicate p-values <0.001 (***) obtained from a Tukey-Kramer multiple comparison test.

Next, we tested whether these proteins co-localize in the same type of membrane nanodomain. For this, we applied high-resolution live cell imaging of root epidermal cells followed by advanced quantitative image analysis of all observations. For evaluating co-localization experiments, we further refined our original quantitative image analysis workflow (3, 4). Here, we compared obtained Pearson correlation coefficients (Rr) to values generated on the corresponding image pairs where blocks of 10x10 pixels in one channel were randomized (randomized Rr= rdRr). Global image analyses combined with image randomizations allow statistically sound statements on the nature of co-localizations (positive, negative, random). Solely expressed SYMREM1 labelled nanodomains in root epidermal cells with a density of 0.077 domains/μm^2^ (Fig. 3H,L). Co-expression of FLOT4 and SYMREM1 in roots in the A17 wild-type background revealed significant degrees of co-localization of both proteins in these cells with an average Pearson correlation coefficient of Rr=0.344 ± 0.027 (Fig. 3A-G) as well as a 4.5-fold increase in SYMREM1-labelled nanodomains (Fig. 3L). Reciprocally, density values for SYMREM1-labelled nanodomains in root hairs were decreased by more than 6-fold when expressing a YFP-SYMREM1 construct in the *flot4-3* mutant background compared to wild-type roots transformed with the same construct (Fig. 3J,K,M). Similar results were also obtained in epidermal cells when expressing a previously published FLOT4-RNAi construct (11) together with the YFP-SYMREM1 using the *Medicago* A17 accession as a genetic background (Fig. 3H,I,N). These data clearly demonstrate that SYMREM1 is recruited into nanodomains in a FLOT4-dependent manner where FLOT4 serves as a central hub during primary nanodomain assembly. Since associations of remorins with the cytoskeleton have been reported (4, 28, 29) we tested whether SYMREM1-recruitment to nanodomains requires an intact cytoskeleton. Pilot experiments where microtubules (Fig. S2A,B) and actin (Fig. S2C,D) were successfully destabilized upon treatment with Oryzalin and Cytochalasin D, respectively, confirmed functionality of the treatment in *M. truncatula* roots. Indeed, nucleation of SYMREM1 in nanodomains appeared to be actin-dependent, since application of Cytochalasin D resulted in a strong and highly significant reduction of SYMREM1-nanodomain density (Fig. S2E,F,H). By contrast, destabilizing microtubules by Oryzalin did not significantly affect SYMREM1 domain patterning (Fig. S2E,G,I).

**Fig. 3.**
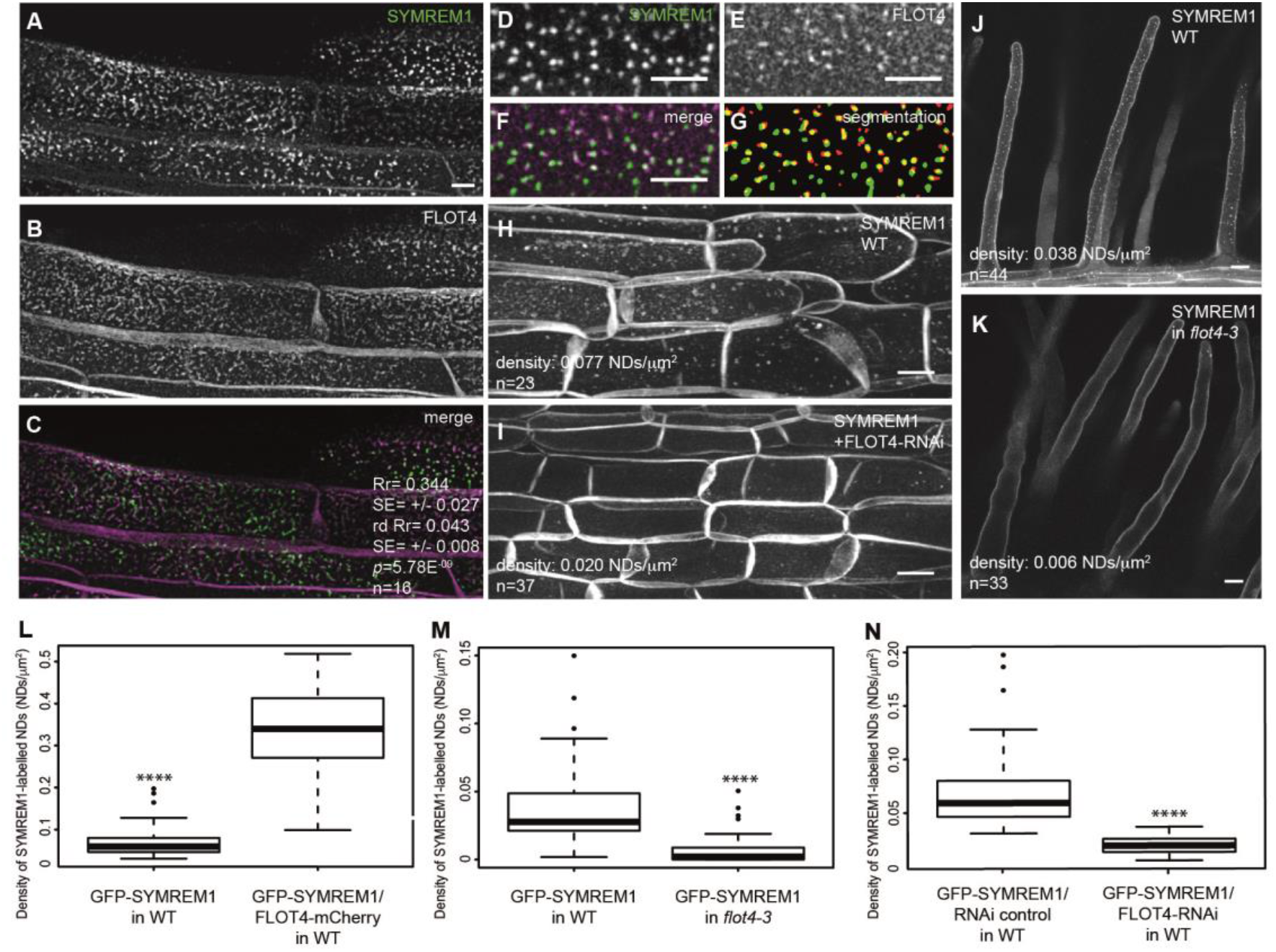
SYMREM1 is recruited to nanodomains in a FLOT4-dependent manner. GFP-SYMREM1 (*A*) and FLOT4-mCherry (*B*) co-localized (*C*) in transgenic *M. truncatula* root epidermal cells. Quantification data are depicted within images. Rr= Pearson correlation coefficient; rd Rr= Pearson correlation coefficient obtained after image randomization of the FLOT4-mCherry channel. SE= standard errors. *p*= confidence interval obtained from a Student t-test comparing Rr and rd Rr with the number of replicates (n) being indicated in the panels. Close-ups of nanodomain-localized GFP-SYMREM1 (*D*) and FLOT4-Cherry (*E*) at the cell surface. Co-localization was quantified in merged images (*F*) and after image segmentation (*G*). Density of GFP-SYMREM1-labelled nanodomains was greatly reduced upon co-expression with a FLOT4-RNAi construct in root epidermal cells (*I*) and in *flot4-3* mutant root hairs (*K*) compared to the respective control (*H,J).* (*L*) Density of GFP-SYMREM1-labelled nanodomains was significantly increased when coexpressed with FLOT4-mCherry. (*M,N*) Quantification on GFP-SYMREM1-labelled nanodomains in the *flot4-3* mutant (*M*) and FLOT4-RNAi roots (*N*). **** indicates *p*-values <0.0001 obtained from student t-tests. Scale bars indicate 5 μm (*A-F*) and 10 μm (*H-K*).

Since SYMREM1 can physically associate with LYK3 and LYK3 immobilization in FLOT4-labelled nanodomains temporally coincides with induced *SYMREM1* expression (6, 26), we tested whether the presence of SYMREM1 at the onset of rhizobial infection of root hairs controls the stimulus-dependent recruitment of LYK3 into these membrane structures. For this, we first verified LYK3 mobility patterns under our laboratory conditions using the same genetic material (*hcl-1* mutant complemented with a ProLYK3:LYK3-GFP construct) and imaging conditions as described earlier (26). Time-lapse imaging of LYK3-GFP driven by its endogenous promoter was performed over 180 seconds and image sections were subjected to kymograph analysis. Vertical band and diffuse patterns indicate locally immobile protein clusters and protein mobility, respectively (Fig. 4). The receptor was indeed mobile in uninoculated roots with pixel dwell times of <30 s (19/20 roots) (Fig. 4A; Table S1). Inoculation of these roots with *S. meliloti* for 16 hours significantly increased dwell times to >180s in 9/9 roots (Fig. 4B; Table S1) confirming ligand-induced immobilization of LYK3 (26). Strikingly, this nanodomain recruitment of LYK3 was also observed when ectopically expressing SYMREM1 in uninoculated roots, i.e. in the absence of endogenous SYMREM1, with a majority of pixel dwell times reaching values of >180s (Fig. 4C; Table S1). Thus, the presence of SYMREM1 is sufficient for the immobilization of LYK3 in nanodomains and independent of any other infection-induced genes. To further elucidate whether presence of SYMREM1 is not only sufficient but also genetically required for the observed mobility arrest of LYK3, we generated transgenic roots in the *symrem1-2* mutant and the R108 WT background expressing the ProLYK3:LYK3-GFP construct. As expected and independently of the genotype, the LYK3 receptor was laterally mobile at root hair plasma membranes in the absence of rhizobia in WT (Fig. 4D) and *symrem1-2* mutant plants (Fig. 4E). However, application of rhizobia to *symrem1-2* mutants resulted in a dramatic change in LYK3 localization (Fig. 4F). Here, the receptor localized to mobile, endosome-like vesicles in 64% of *symrem1-2* roots while similar structure were only observed in 24% of the control roots (Fig. 4F-H’’’,I). Plasma membrane origin of these structures was confirmed by staining membranes with the lipophilic styryl dye FM4-64 (Fig. S3). Interestingly, this effect was not observed when conducting the same experiment in the *flot4-3* mutant background (Fig. 4I) albeit endosomal structures were more frequently in *flot4-3* (26%) compared to wild-type roots (13%) in the absence of rhizobia. Different to *symrem1-2* only a modest increase in the frequency of LYK3 endocytosis was observed in *flot4-3* mutants and WT plants upon inoculation with rhizobia (Fig. 4I). This indicates that infection-triggered endocytosis of LYK3 is controlled by SYMREM1 rather than FLOT4. These data unambiguously show that, in contrast to a possible LYK3 endocytosis that is observed 6 hours after NF application (and in the absence of rhizobia and endogenous SYMREM1) (26), infection-induced expression and accumulation of the molecular scaffold protein SYMREM1 mediates stabilization of the activated receptor pool by receptor recruitment to membrane nanodomains. Consequently, SYMREM1 acts as a negative regulator of receptor endocytosis, a function that is required for successful infection as implied by the *symrem1* mutant phenotype. Considering the overlapping IT phenotypes in *LYK3* knockdown experiments (23) and *symrem1* mutants (Fig. 2) it can be concluded that IT abortion in *symrem1* mutants is caused by a depletion of LYK3 from IT membranes. Taken together our data genetically prove that scaffold-mediated recruitment of LYK3 to specific nanodomains is mandatory for root hair infection. This in turn may provide the host with a mechanism that allows spatio-temporal desensitisation of cell surfaces towards the respective ligand as the host might not be able to durably stabilize the activated NFP/LYK3 receptor complex at the plasma membrane. Prematurely aborted infections as observed in *flot4, symrem1* and *lyk3* mutants indicate that stabilization of LYK3 during host cell penetration is required for controlling cellular entry along the root and during IT guidance towards the root cortex.

**Fig. 4.**
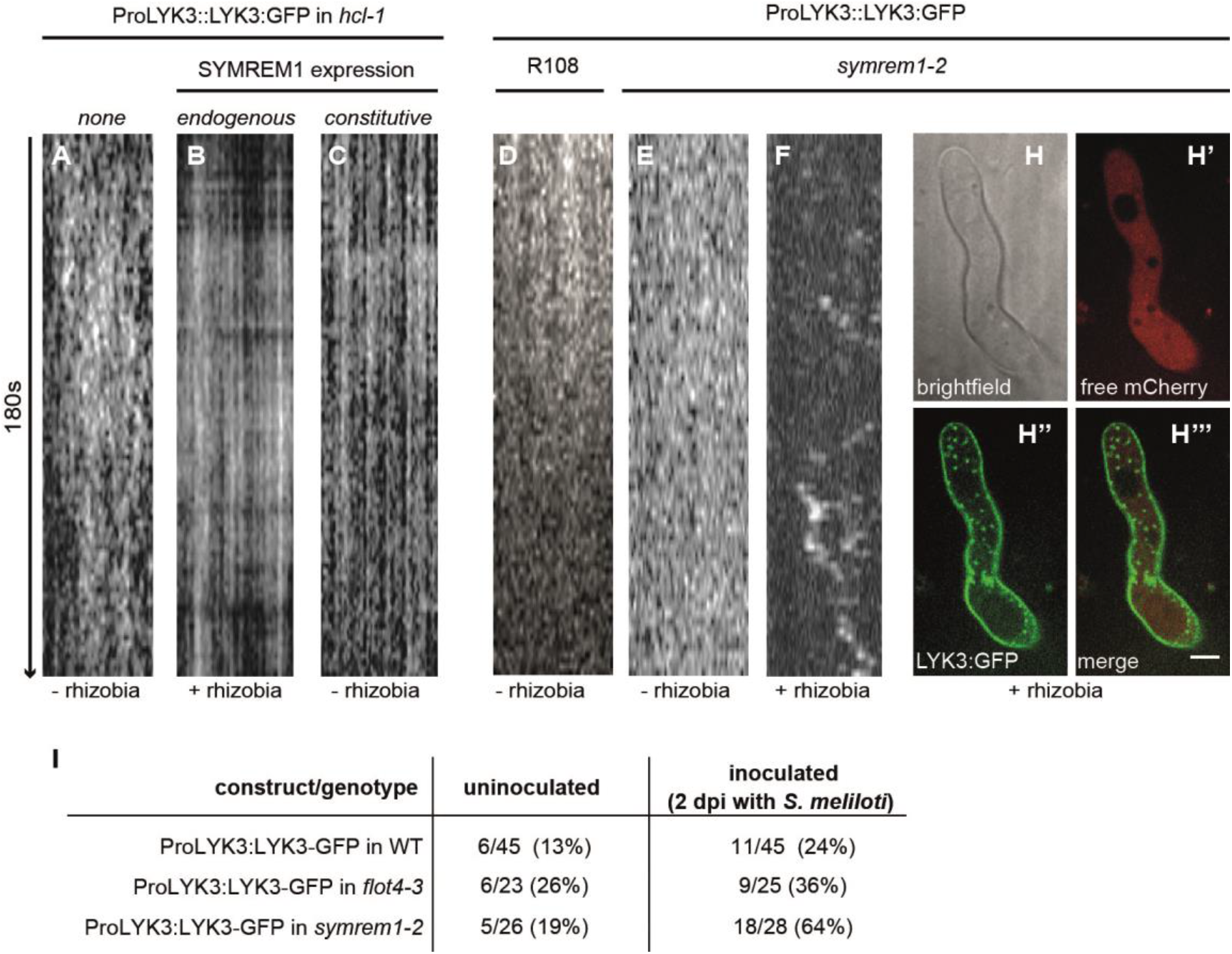
SYMREM1 negatively regulates infection-triggered receptor endocytosis by stabilizing LYK3 in nanodomains. Kymograph analyses of time-lapse image series taken from cell surfaces at the base of *M. truncatula* root hairs. LYK3 is mobile in the absence of rhizobia (*A*) and gets immobilized 16 hours after rhizobial inoculation, i.e. at the onset of infection (*B*). The same effect was obtained when ectopically expressing SYMREM1 in the absence of rhizobia (i.e. in the absence of endogenous SYMREM1) (*C).* Analysis of uninoculated transgenic *M. truncatula* roots expressing LYK3-GFP under its native promoter revealed receptor mobility in WT R108 (*D*) and the *symrem1-2* (*E*) mutant background. Inoculation of LYK3-GFP expressing *symrem1-2* mutant roots, resulted in destabilization of the receptor (*F*) and its accumulation in endosome-like vesicles (H-H’’’). Scale bars indicate 10 μm. (*I*) Frequency of LYK3 endocytosis in different genotypes (R108 WT, *symrem1-2, flot4-3*) was scored in independent root hairs under uninoculated conditions and 2 dpi with. *S. meliloti.*

## MATERIALS AND METHODS

### Plant growth and phenotypical analysis

For phenotypical analysis *Medicago truncatula* wild-type R108, *symrem1-1* (NF4432), *symrem1-2* (NF4395) and *flot4-3* (NF14107) seeds were scarified and sterilized before being sown on 1% agar plates for germination and kept in dark at 4°C for 3-5 days for vernalization. Germination was allowed for up to 24 hours at 24°C before transferring the seedlings to Fahraeus plates. Plants were grown for 4 days in the presence of 1 μM nitrate before being transferred to plates without nitrogen but containing 0.1 μM aminoethoxyvinylglycine (AVG) for 3 days. Plants were then inoculated with 1ml *Sinorhizobium meliloti* 2011 (lacZ) with an OD600 of 0.05. Infection threads were scored 7 dpi.

### Genotyping of Tnt1 Insertion Lines and Quantitative Real-Time PCR

R0 or R1 seeds of *M. truncatula* R108 *Tnt1* transposon insertion lines were obtained from the Samuel Roberts Noble Foundation after screening the mutant population by PCR using the primers described in Table S2.

Total RNA of control and insertion lines was extracted using a commercial kit (Promega) following the supplier’s instructions. An additional DNaseI treatment (Invitrogen) was performed. Synthesis of cDNA and qRT-PCR were conducted as described in earlier using the SuperScriptIII reverse Transcriptase (Invitrogen). All data were normalized to Ct values of the housekeeping gene ubiquitin (30), the related primers are described in Table S2.

### Hairy Root Transformation and inoculation of rhizobia

*M. truncatula* hairy root transformation was performed as previously described (31) using the *Agrobacterium rhizogenes* strain ARqua1 and transferred weekly to fresh plates containing Fahraeus medium (pH 6.0) supplemented with 1 mM NH_4_NO_3_ and followed by 2 days of growth in nitrogen-free Fahraeus medium containing 0.1 μM AVG. Images for kymograph and immune-fluorescence analyses were taken on root hairs on plants inoculated for two and six days, respectively.

### Construct Design

The coding sequence of *Medicago truncatula* SYMREM1 (Genbank accession JQ061257) was recombined into the Gateway (GW) compatible ProUBi-YFP-GW vector (12) via LR-reaction. For complementation experiments a Gateway compatible ProSYMREM1-GW vector was created by replacing the Ubiquitin promoter with the functional SYMREM1 promoter (643bp upstream of the translational start of *SYMREM1*) using PmeI and XbaI restriction sites. All other constructs were cloned as Golden Gate compatible constructs. BpiI and BsaI restriction sites were removed from the following nucleotide templates prior to the cloning of Level 2 expression vectors: *SYMREM1* (Genbank accession: JQ061257), *MAP4* (Genbank accession: M72414), the cDNA of the genomic *FLOT4* (Genbank accession GU224281), the 2kb *FLOT4* promoter (ProFLOT4), as well as the *SYMREM1* promoter (ProSYMREM1). A Lifeact template with flanking BsaI restriction sites were directly inserted into pUC-Bpi via blunt end StuI (NEB) cut-ligation for subsequent Golden Gate cloning. Double stranded sequences for the FLOT4-RNAi constructs with flanking BsaI sites were also cloned via a blunt end StuI cut-ligation into pUC-BpiI. RNAi silencing vectors based on sequenced described earlier (11) were assembled as previously described (32). For localization of FLOT4 and SYMREM1 driven by endogenous promoters, a tandem cassette ProUbi:NLS-2xCerulean was inserted as the marker for selecting transgenic roots prior to analysis. All the designed constructs are listed in Table S2.

### Immunolocalization assay

An improved procedure for whole-mount immune-localization was adopted (33). In brief, 0.5 - 1 cm root pieces were mounted in 2 *%* formaldehyde in MTSB buffer supplemented with 0.1 % Triton. Vacuum infiltration of explants, cuticle solubilisation, cell walls digestion and membrane permeabilisation were performed as originally described (33). After 1 hour blocking in BSA samples were incubated with a GFP nano-booster (1:200 dillution) coupled to Atto488 (Chromotek, Germany) for 1 hour at room temperature followed by three rounds of 10 min washing before being mounted in anti-fade medium.

### Confocal Laser-Scanning Microscopy

For cytoskeleton and NLS-GFP reporter images, the confocal laser scanning microscopy was performed on a Leica TCS SP5 confocal microscope equipped with 63x and 20x HCX PL APO water immersion lenses (Leica Microsystems, Mannheim, Germany). GFP was excited with the Argon laser (AR) line at 488 nm and the emission detected at 500-550 nm. YFP was excited with the 514 nm AR laser line and detected at 520-555 nm. mCherry fluorescence was excited using the Diode Pumped Solid State (DPSS) laser at 561nm and emission was detected between 575-630 nm. Samples, co-expressing two fluorophores were imaged in sequential mode between frames. Due to low signal intensity for the ProSYMREM1-NLS-2xGFP reporter, the corresponding fluorescence was detected using Leica HyD detectors. Images were taken with a Leica DFC350FX digital camera.

Nanodomain patterns for FLOT4-GFP, YFP-SYMREM1 and GFP-SYMREM1 were acquired using a Leica TCS SP8 confocal microscope system with 63x HCX PL APO CS2 oil immersion and 20x HCX PL APO water immersion lenses. YFP was excited with the 514 nm AR laser line and detected at 520–555nm. GFP was excited by White Light laser at 488 nm and emission was detected between 500-550 nm, the corresponding fluorescence was detected using Leica HyD detectors. mCherry fluorescence from *Sinorhizobium meliloti* was excited using White Light laser at 561 nm and emission was detected between 575-630 nm. To create z-stacks of 35-50 μm depth images were taken with 0.5-1 μm space intervals.

LYK3 mobility was assessed using a Perkin Elmer UltraView Vox on a inverted Zeiss microscope stand equipped with a Yokogawa spinning disk unit and a Hamamatsu EMCCD C9100-50 camera. GFP was excited with a 488 nm laser set to 30% with an exposure time of 900 ms. Camera binning was set to 2 during images acquisition. Time series data was collected over 90 frames taken at 2-second intervals, for a total of 180 seconds. The endocytosis with FM4-64 staining was imaged using Zeiss LSM 880 with Airyscan in the Fast Airyscan mode. GFP and FM4-64 were excited with 488 nm and 561 nm, respectively.

### Quantitative Image Analysis

Image analysis was performed with the open source ImageJ/ (Fiji) software (34). For illustration, images were background subtracted according to the rolling ball algorithm, filtered with a Mean filter pixel radius of 1 and then maximum z-projected (create stack). Contrast was enhanced for visualization in Figures but not for quantification. Pixel based co-localizations to determine Pearson Correlation Coefficient values were performed using the Fiji Plugins ‘Squassh’(35) and ‘JACoP’ (36). Image segmentation was performed with ‘Squassh’. Randomization was performed with the automatic Costes’ Randomization method in ‘JACoP’ in which clusters of 10x10 pixels were randomly distributed in one channel and correlated to the original values. Additionally, randomization was also performed on maximum z-projections via horizontal flip of the mCherry channel as described previously (4, 37). To quantify nanodomains images were segmented to differentiate background from domains. For this, the background was subtracted using a rolling ball algorithm with a radius corresponding to the largest structure of interest. A mean blur with radius 1 was then applied, and the slices (n=5-12 slices, with distances of 0.25 to 0.7 μm) maximum projected along the z-axis for epidermal cells and the slice with distances of 0.5 μm was used for root hair images. A threshold was applied to the images and the result saved as a binary mask. The ‘create selection’ tool was used to mark the outlines and was overlaid onto the original image to verify proper image segmentation. Domains were counted with the ‘particle analyzer’ tool in Fiji. For mobility analysis of LYK3-GFP nanodomain, kymographs were generated using ‘‘Reslice’’ tool in Fiji.

### Cytoskeleton depolymerisation

A 1mM Oryzalin stock solution in DMSO and a 10mM Cytochalasin D stock solution in EtOH were prepared. *Medicago truncatula* root samples of 1 cm length were incubated in final concentrations of 10 μM Oryzalin or 10 μM Cytochalasin D for 12 hours in water. The control samples were incubated in water with the equal amount of solvent for the same time.

## ACKNOWLEDGEMENTS

We would like to thank all members of our team for their fruitful discussions throughout the project and comments on the manuscript. We would also like to acknowledge Cara Haney (University of British Columbia, CA) for sharing some unpublished experimental details. The complemented *hcl-1* line and the ProLYK3:LYK3-GFP construct were kindly provided by Doug Cook and Brendan Riely (UC Davis, USA). We thank the staff of the Life Imaging Center (LIC) in the Center for Biological Systems Analysis (ZBSA) of the Albert-Ludwigs-University Freiburg and the Center for Advanced Light Microscopy (CALM) at the University of Munich for help with their confocal microscopy resources, and the excellent support in image recording. We also greatly appreciate the methodological input of Taras Pasternak (University of Freiburg, Germany). The *Medicago truncatula* plants utilized in this research project, which are jointly owned by the Centre National De La Recherche Scientifique, were obtained from The Samuel Roberts Noble Foundation, Inc. and were created through research funded, in part, by a grant from the National Science Foundation, NSF# 703285. This work was funded by an Emmy-Noether grant of the Deutsche Forschungsgemeinschaft (DFG; OT423/2-1; TO) and the China Scholarship Council (CSC) (grant no. 201506350004 to PL).

## SUPPORTING INFORMATION

**Table S1:** Quantification of LYK3 mobility. Pixel dwell times of LYK3-labelled nanodomains were categorized as >180 seconds (s), 60-180 seconds and <30 seconds. Values describe frequencies of categories detected in a set of observed root hairs (n=9-49).

| Mobility | uninoculated | *S. meliloti* (2 dpi) | ProUbi:YFP-SYMREM1 |
| --- | --- | --- | --- |
| >180s | 1/20 | 9/9 | 30/49 |
| 60-180s |  |  | 16/49 |
| <30s | 19/20 |  | 3/49 |

## SUPPORTING INFORMATION - FIGURES

**Fig. S1.**
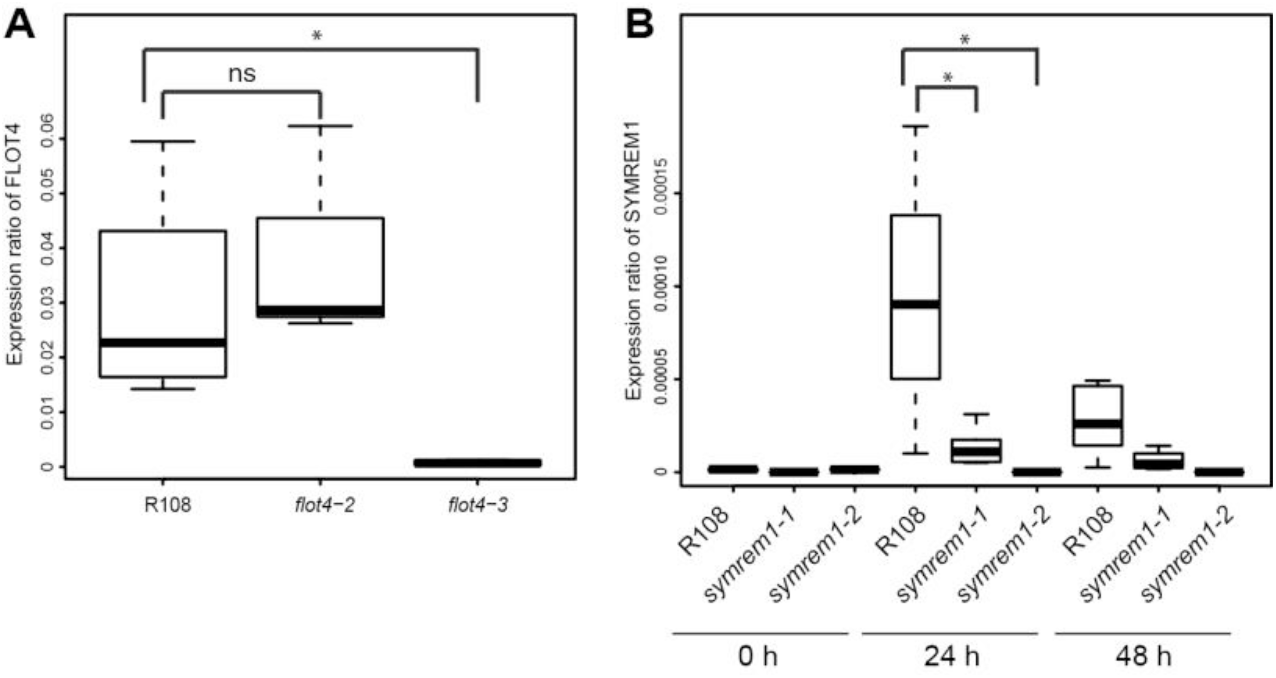
Determining transcript and protein levels in *Tnt1* insertion lines. (*A*) *FLOT4* transcript levels in *flot4-2* and *flot4-3* insertion lines and wild-type R108 plants. (*B*) *SYMREM1* transcript level before inoculation, at 24 hour and 48 hours post inoculation, in *symrem1-1* and *symrem1-2* mutant lines. Quantitative qRT-PCR was performed on cDNA obtained from roots, with 5 biological replicates. The graphs represent the ΔΔCt values obtained by qRT-PCR in relative to ubiquitin. * indicates *p*-values <0.05 obtained from student t-tests in A, and from Dunnett multiple comparison test in B, ‘ns’= not significant.

**Fig. S2.**
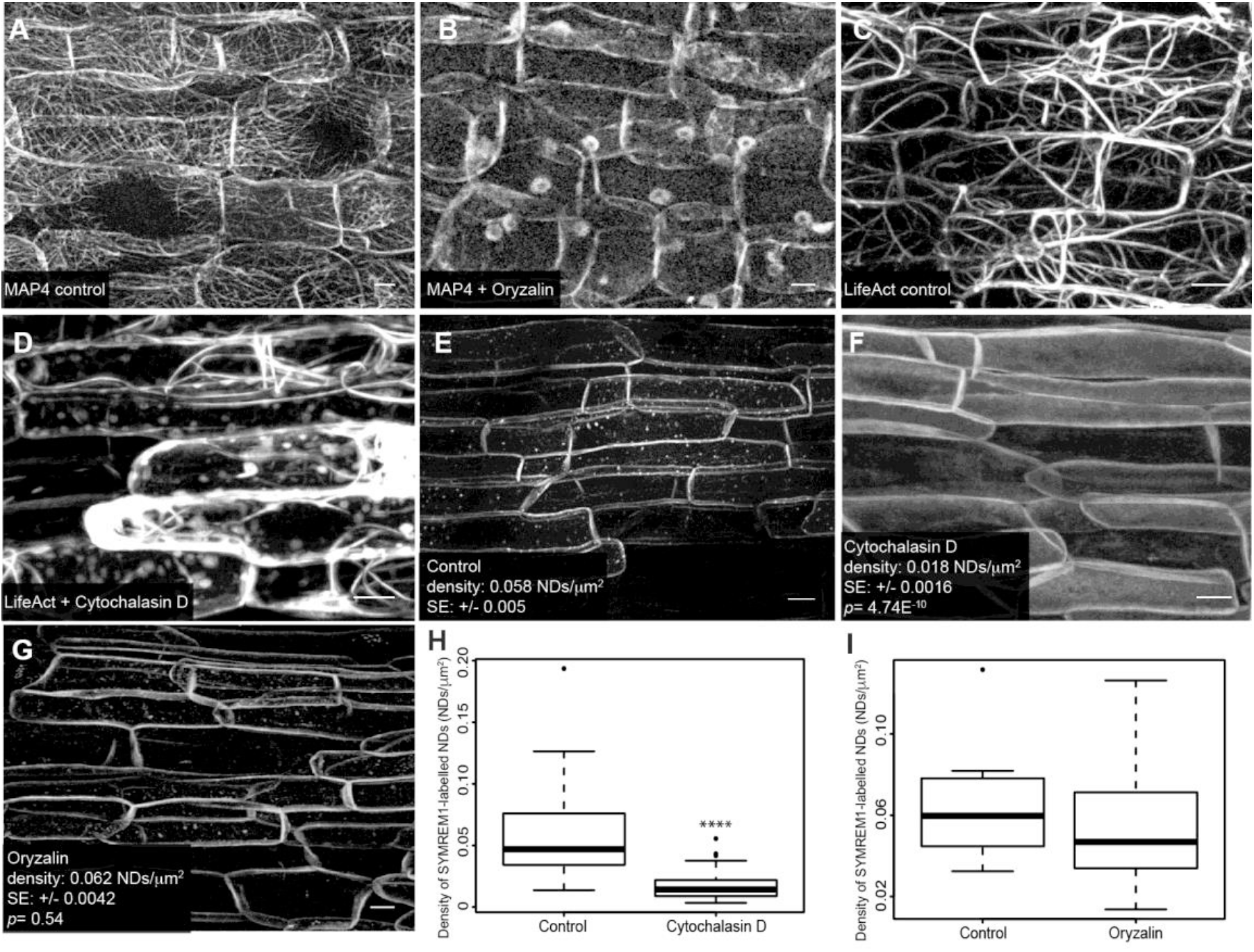
Recruitment of SYMREM1 into nanodomains is actin-dependent. (*A-D*) Efficiency of drug application to destabilize microtubules (*A,B*) and actin (*D,E*) was tested by applying Oryzalin (*B*) and Cytochalasin D (*D*) on roots expressing fluorophore-tagged MICROTUBULE-ASSOCIATED PROTEIN 4 (MAP4) (A,B) or LifeAct (C,D), respectively. (*E-I*) Density of YFP-SYMREM1-labelled nanodomains was significantly higher in control roots (*E,H*) compared to those treated with Cytochalasin D (*F, H*) while depolymerising microtubules (*G,I*) did not change YFP-SYMREM1 localization. Quantitative image analysis was performed on all samples as indicated below the individual panels. SE= standard error; **** indicate *p*-values <0.0001 obtained from student t-tests. Scale bars indicate 10 μm.

**Fig. S3.**
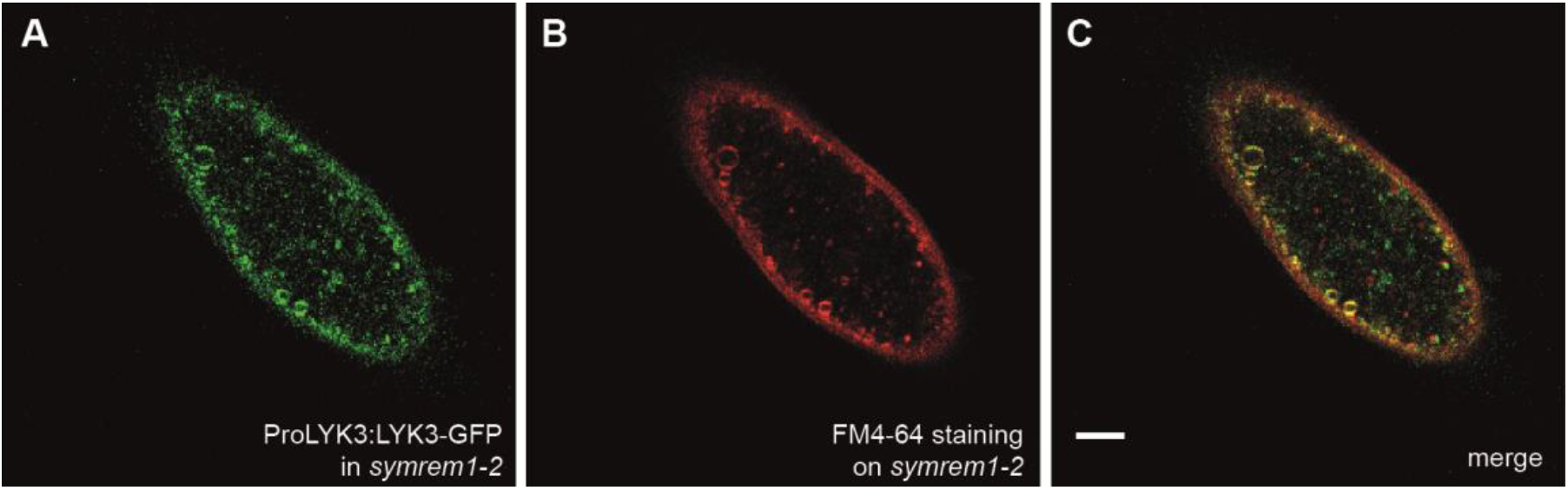
Activated LYK3 receptor localizes to endosomes in *symrem1-2* mutants upon prolonged rhizobial inoculation. ProLYK3:LYK3-GFP was expressed in *symrem1-2* mutant roots and imaged in root hairs 2 dpi with *S. meliloti* (*A*). To test PM origin of the observed vesicles samples were counterstained with FM4-64 (*B*) and images were subsequently merged (*C*). (*D*) Fluorescence intensity plot of a representative transect (position indicated in (*C*)) shows high degrees of co-localization in these structures. Scale bar indicates 5 μm.

**Fig. S4.**
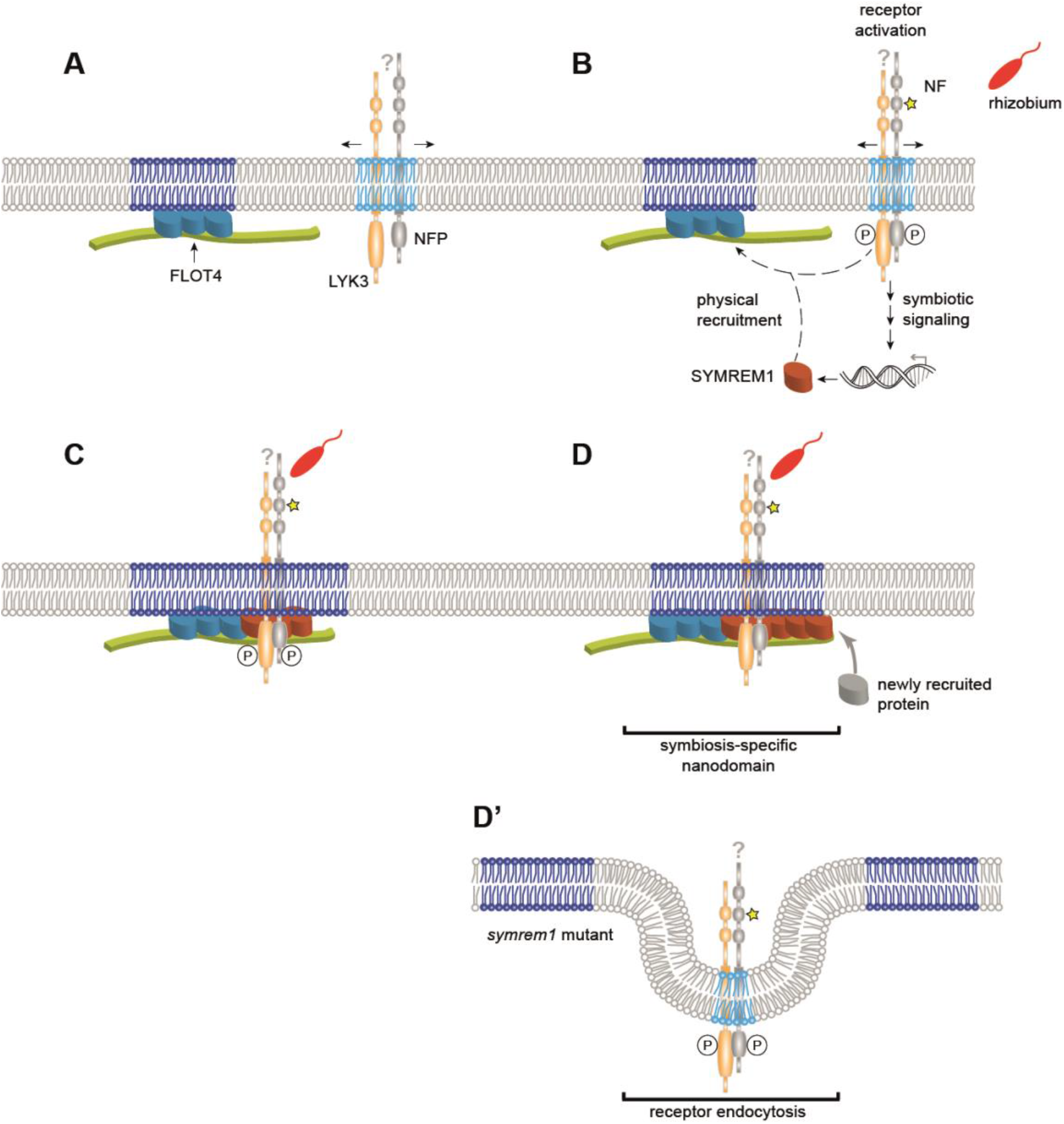
Proposed model for nanodomain assembly. (*A*) Constitutively expressed FLOT4 (turquoise) forms a primary nanodomain scaffold that is unable to recruit LYK3 in the absence of SYMREM1. (*B*) Nod factor (NF) perception by NFP (grey) and LYK3 (orange) occurs in mobile nanodomains and results in the activation of a symbiosis-specific signalling cascade that leads to the expression of *SYMREM1* (red). (*C*) Due to its ability to directly bind LYK3 (6), SYMREM1 actively recruits the receptor into the FLOT4 nanodomain. (*D*) Phosphorylation of SYMREM1 by LYK3 might trigger remorin oligomerization, which generates new docking sites for proteins required for rhizobial infection (hypothetical). (*D’*) In *symrem1* mutants LYK3 is destabilized and endocytosed upon rhizobial inoculation.

## REFERENCES

1. Burkart RC & Stahl Y (2017) Dynamic complexity: plant receptor complexes at the plasma membrane. Curr Opin Plant Biol 40:15–21.

2. Ott T (2017) Membrane nanodomains and microdomains in plant-microbe interactions. Curr Opin Plant Biol 40:82–88.

3. Bücherl CA, et al. (2017) Plant immune and growth receptors share common signalling components but localise to distinct plasma membrane nanodomains. eLife 6:e25114.

4. Jarsch IK, et al. (2014) Plasma Membranes Are Subcompartmentalized into a Plethora of Coexisting and Diverse Microdomains in Arabidopsis and Nicotiana benthamiana. Plant Cell 26(4):1698–1711.

5. Gui J, et al. (2016) OsREM4.1 Interacts with OsSERK1 to Coordinate the Interlinking between Abscisic Acid and Brassinosteroid Signaling in Rice. Dev Cell 38(2):201–213.

6. Lefebvre B, et al. (2010) A remorin protein interacts with symbiotic receptors and regulates bacterial infection. Proc Natl Acad Sci U S A 107(5):2343–2348.

7. Tóth K, et al. (2012) Functional Domain Analysis of the Remorin Protein LjSYMREM1 in Lotus japonicus. PLOS ONE 7(1):e30817.

8. Wang X, et al. (2015) Single-molecule fluorescence imaging to quantify membrane protein dynamics and oligomerization in living plant cells. Nat Protoc 10(12):2054–2063.

9. Jarsch IK & Ott T (2011) Perspectives on remorin proteins, membrane rafts, and their role during plant-microbe interactions. Mol Plant Microbe Interact 24(1):7–12.

10. Gronnier J, et al. (2017) Structural basis for plant plasma membrane protein dynamics and organization into functional nanodomains. eLife 6:e26404.

11. Haney CH & Long SR (2010) Plant flotillins are required for infection by nitrogen-fixing bacteria. Proc Natl Acad Sci U S A 107(1):478–483.

12. Konrad SS, et al. (2014) S-acylation anchors remorin proteins to the plasma membrane but does not primarily determine their localization in membrane microdomains. New Phytol 203(3):758–769.

13. Li R, et al. (2012) A membrane microdomain-associated protein, Arabidopsis Flot1, is involved in a clathrin-independent endocytic pathway and is required for seedling development. Plant Cell 24(5):2105–2122.

14. Perraki A, et al. (2012) Plasma Membrane Localization of Solanum tuberosum Remorin from Group 1, Homolog 3 Is Mediated by Conformational Changes in a Novel C-Terminal Anchor and Required for the Restriction of Potato Virus X Movement]. Plant Physiol 160(2):624–637.

15. Raffaele S, et al. (2009) Remorin, a Solanaceae protein resident in membrane rafts and plasmodesmata, impairs potato virus X movement. Plant Cell 21(5):1541–1555.

16. Gui J, Liu C, Shen J, & Li L (2014) Grain setting defect1, encoding a remorin protein, affects the grain setting in rice through regulating plasmodesmatal conductance. Plant Physiol 166(3):1463–1478.

17. Checker VG & Khurana P (2013) Molecular and functional characterization of mulberry EST encoding remorin (MiREM) involved in abiotic stress. Plant Cell Rep 32(11):1729–1741.

18. Yue J, Li C, Liu Y, & Yu J (2014) A remorin gene SiREM6, the target gene of SiARDP, from foxtail millet (Setaria italica) promotes high salt tolerance in transgenic Arabidopsis. PLOS ONE 9(6):e100772.

19. Perraki A, et al. (2014) StRemorin1.3 hampers Potato virus X TGBp1 ability to increase plasmodesmata permeability, but does not interfere with its silencing suppressor activity. FEBS Lett 588(9):1699–1705.

20. Son S, Oh CJ, Bae JH, Lee H, & An CS (2015) GmREM1.1 and GmREM2.1, Which Encode the Remorin Proteins in Soybean, Have Distinct Roles during Root Nodule Development. J Plant Biol 58:17–25.

21. Zipfel C & Oldroyd GE (2017) Plant signalling in symbiosis and immunity. Nature 543(7645):328–336.

22. Limpens E, et al. (2003) LysM domain receptor kinases regulating rhizobial Nod factor-induced infection. Science 302(5645):630–633.

23. Smit P, et al. (2007) Medicago LYK3, an entry receptor in rhizobial nodulation factor signaling. Plant Physiol 145(1):183–191.

24. Murray JD (2011) Invasion by invitation: rhizobial infection in legumes. Mol Plant Microbe Interact 24(6):631–639.

25. Fournier J, et al. (2015) Remodeling of the infection chamber before infection thread formation reveals a two-step mechanism for rhizobial entry into the host legume root hair. Plant Physiol 167(4):1233–1242.

26. Haney CH, et al. (2011) Symbiotic Rhizobia Bacteria Trigger a Change in Localization and Dynamics of the Medicago truncatula Receptor Kinase LYK3. Plant Cell 23(7):2774–2787.

27. Tadege M, et al. (2008) Large-scale insertional mutagenesis using the Tnt1 retrotransposon in the model legume Medicago truncatula. Plant J 54(2):335–347.

28. Gui J, Zheng S, Shen J, & Li L (2015) Grain setting defect1 (GSD1) function in rice depends on S-acylation and interacts with actin 1 (OsACT1) at its C-terminal. Front Plant Sci 6:804.

29. Szymanski WG, et al. (2015) Cytoskeletal Components Define Protein Location to Membrane Microdomains. Mol Cell Proteomics 14(9):2493–2509.

30. Satge C, et al. (2016) Reprogramming of DNA methylation is critical for nodule development in Medicago truncatula. Nat Plants 2(11):16166.

31. Boisson-Dernier A, et al. (2001) Agrobacterium rhizogenes-transformed roots of Medicago truncatula for the study of nitrogen-fixing and endomycorrhizal symbiotic associations. Mol Plant Microbe Interact 14(6):695–700.

32. Binder A, et al. (2014) A modular plasmid assembly kit for multigene expression, gene silencing and silencing rescue in plants. PLOS ONE 9(2):e88218.

33. Pasternak T, et al. (2015) Protocol: an improved and universal procedure for whole-mount immunolocalization in plants. Plant Methods 11:50.

34. Schindelin J, et al. (2012) Fiji: an open-source platform for biological-image analysis. Nat Methods 9(7):676–682.

35. Rizk A, et al. (2014) Segmentation and quantification of subcellular structures in fluorescence microscopy images using Squassh. Nat Protoc 9(3):586–596.

36. Bolte S & Cordelieres FP (2006) A guided tour into subcellular colocalization analysis in light microscopy. JMicrosc 224(Pt 3):213–232.

37. Jarsch IK & Ott T (2015) Quantitative Image Analysis of Membrane Microdomains Labelled by Fluorescently Tagged Proteins in Arabidopsis thaliana and Nicotiana benthamiana. bio-protocol 5(11):e1497.

